# Streptavidins play a multifunctional role within a biotin-pathway antibiotic network encoded in a biosynthetic supercluster

**DOI:** 10.1101/2025.11.10.687665

**Authors:** Sumire Kurosawa, Jingjun Mo, Xiaotong Diao, Feng Xie, Tingting Wang, Haowen Zhao, Dalei Wu, Rolf Müller, Chengzhang Fu

**Affiliations:** Helmholtz Institute for Pharmaceutical Research Saarland (HIPS), Helmholtz Centre for Infection Research (HZI), 66123 Saarbrücken, Germany; Helmholtz International Lab for Anti-Infectives, Helmholtz Center for Infection Research, 38124 Braunschweig, Germany; Helmholtz International Lab for Anti-Infectives, State Key Laboratory of Microbial Technology, Shandong University, Qingdao 266237, China; German Centre for Infection Research (DZIF), 38124 Braunschweig, Germany; PharmaScienceHub, Saarbrücken 66123, Germany

## Abstract

Streptavidin, the well-known biotin-binding protein extensively used in biotechnology, is naturally co-produced in *Streptomyces avidinii* alongside stravidins—inhibitors of biotin biosynthesis. Here, we uncover and activate a conserved genomic region flanked by two streptavidin genes, revealing multiple biosynthetic gene clusters that produce diverse biotin-related metabolites, including stravidins, acidomycin, α-methyl-KAPA, α-methyldesthiobiotin, and the novel non-proteinogenic amino acid ANDA. We show α-methyldesthiobiotin arises from α-methyl-KAPA, illustrating how methylated analogues interfere with distinct steps of biotin biosynthesis. Contrary to the view that streptavidin functions solely by sequestering biotin, our biochemical, structural, and bioactivity analyses demonstrate that it also binds acidomycin, neutralizing its antibacterial activity to protect the producer while likely facilitating compound secretion. The crystal structure of the streptavidin–acidomycin complex reveals the molecular basis for this dual functionality. Our findings establish a multifunctional streptavidin that integrates biotin sequestration and self-resistance to balance an antibiotic network targeting the biotin pathway in microbial competition.

## Introduction

Biotin (vitamin B_7_) is an essential cofactor for carboxylation, decarboxylation, and transcarboxylation reactions that underpin central metabolism, including fatty acid biosynthesis and gluconeogenesis. Unlike many bacteria, humans and other higher organisms cannot synthesize biotin *de novo* and must obtain it from diet or commensal microbiota^1^. Despite its ubiquity, free biotin levels in human plasma are relatively low^2^, suggesting limited availability in host environments. This metabolic asymmetry, where bacteria depend on endogenous biosynthesis while humans rely on external sources, renders the biotin biosynthetic pathway an attractive and selective target for antibacterial drug development^1–3^.

In *Escherichia coli*, the biotin biosynthetic genes form an operon comprising *bioA*, *bioB*, *bioF*, *bioC*, and *bioD*^4–6^. Although pathway organization varies among bacterial species, both *E. coli* and mycobacteria initiate biotin synthesis by constructing the C7 carboxyl moiety through fatty acid biosynthetic enzymes^1,3,7,8^. This precursor is condensed with alanine by BioF to yield 7-keto-8-aminopelargonic acid (KAPA), which is subsequently converted by BioA, BioD, and BioB into 7,8-diaminopelargonic acid (DAPA), desthiobiotin, and finally biotin^1^ (Fig. 1a). During the second half of the 20th century, several biotin-related natural products were discovered, including the BioB inhibitor acidomycin (**1**)^9–13^, the BioA inhibitor stravidin (**2**)^14–16^, α-methyldesthiobiotin/α-methylbiotin (**3/4**)^17–19^, and α-methyl-KAPA (**5**)^20^, although their biosynthetic origin remained elusive for decades (Fig. 1b).

**Fig. 1:**
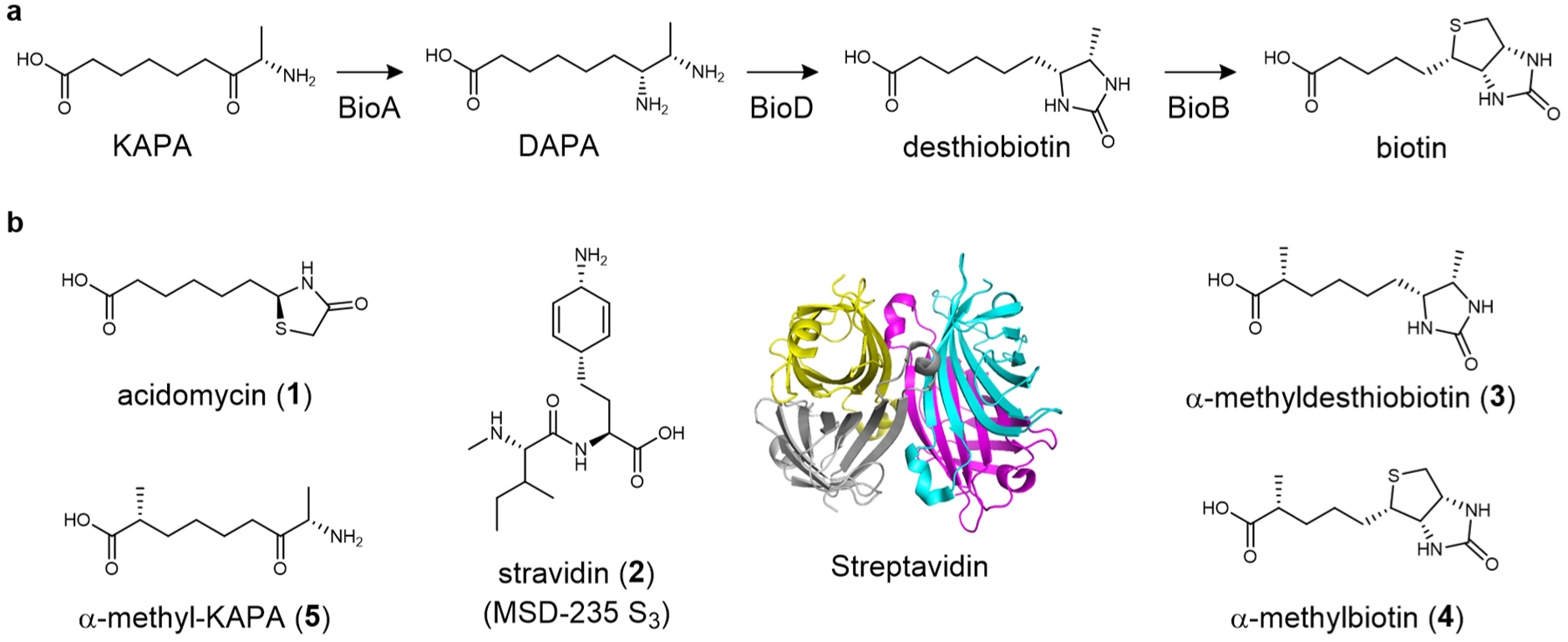
Biotin biosynthesis and known biotin-related compounds/protein. **a**, Biotin biosynthesis. **b**, Known biotin-related compounds and streptavidin (PDB ID: 1STP)^22^.

Stravidins were discovered together with the biotin-binding protein streptavidin from *Streptomyces avidinii*, which became indispensable in biotechnology due to its exceptional affinity for biotin^15,21^. The biological rationale for streptavidin production, however, has remained unclear. It has been proposed that streptavidin sequesters environmental biotin to inhibit competitors, but bacterial capacity for *de novo* biotin biosynthesis challenges this view. The observed synergistic inhibition of *E. coli* by streptavidin and stravidins hints at a more intricate interplay between biotin sequestration and biosynthetic inhibition (Fig. 1a)^15^.

Building on this reasoning, we hypothesized that the streptavidin gene may serve as a genomic anchor for neighboring biosynthetic gene clusters (BGCs) encoding biotin-pathway inhibitors. Here, we uncovered a streptavidin-flanked genomic architecture in *Streptomyces*, a “BGC Nexus” encoding multiple BGCs for biotin-pathway inhibitors (Fig. 1b). Activation of this region using the CRISPR– Cas9-based ACTIMOT platform^23^ revealed a coherent metabolic network producing stravidins, acidomycin, α-methyl-KAPA/α-methyldesthiobiotin, and the previously unknown amino acid ANDA. Individual BGCs were further characterized via heterologous expression, gene deletion, biochemical assays, and comprehensive structure elucidation, including absolute stereochemical determination, providing insight into their biosynthesis, including the conversion of α-methyl-KAPA to α-methyldesthiobiotin.

Beyond investigating these biosynthetic pathways, we uncovered a previously unrecognized role for streptavidin. Structural biology and bioactivity analyses show that, in addition to binding biotin, streptavidin binds acidomycin and neutralizes its antibacterial activity, mediating both self-protection and environmental competition. These findings reveal a streptavidin-anchored genomic system that integrates protein–metabolite interactions to control biotin metabolism and potentially influence microbial interactions.

## Results

### Dual streptavidin genes anchor a genomic region enriched in biotin-related BGCs

The co-discovery and reported synergy of stravidins and streptavidin in *S. avidinii* DSM40526 suggested that their biosynthetic genes might be genomically linked^15^. We therefore set out to locate the stravidin BGC and hypothesized that it would be co-localized with the streptavidin gene, potentially together with other biotin-antimetabolite BGCs. Using the streptavidin gene as a genomic marker, we thus systematically examined the genomes of known producers of biotin-pathway inhibitors.

In the genome of *S. avidinii* DSM40526, we indeed identified a second streptavidin gene (*sav2*), located ∼36 kb away from the canonical *sav1* and ∼25 kb from the primary biotin operon (Fig. 2a). Between *sav1* and *sav2* lies a compact operon, *staA–staN*, predicted to encode the stravidin BGC based on its enzymatic repertoire for chorismate and branched-chain amino acid metabolism^24^. This assignment is consistent with a recent report describing the same BGC from another *Streptomyces* strain^25^.

**Fig. 2:**
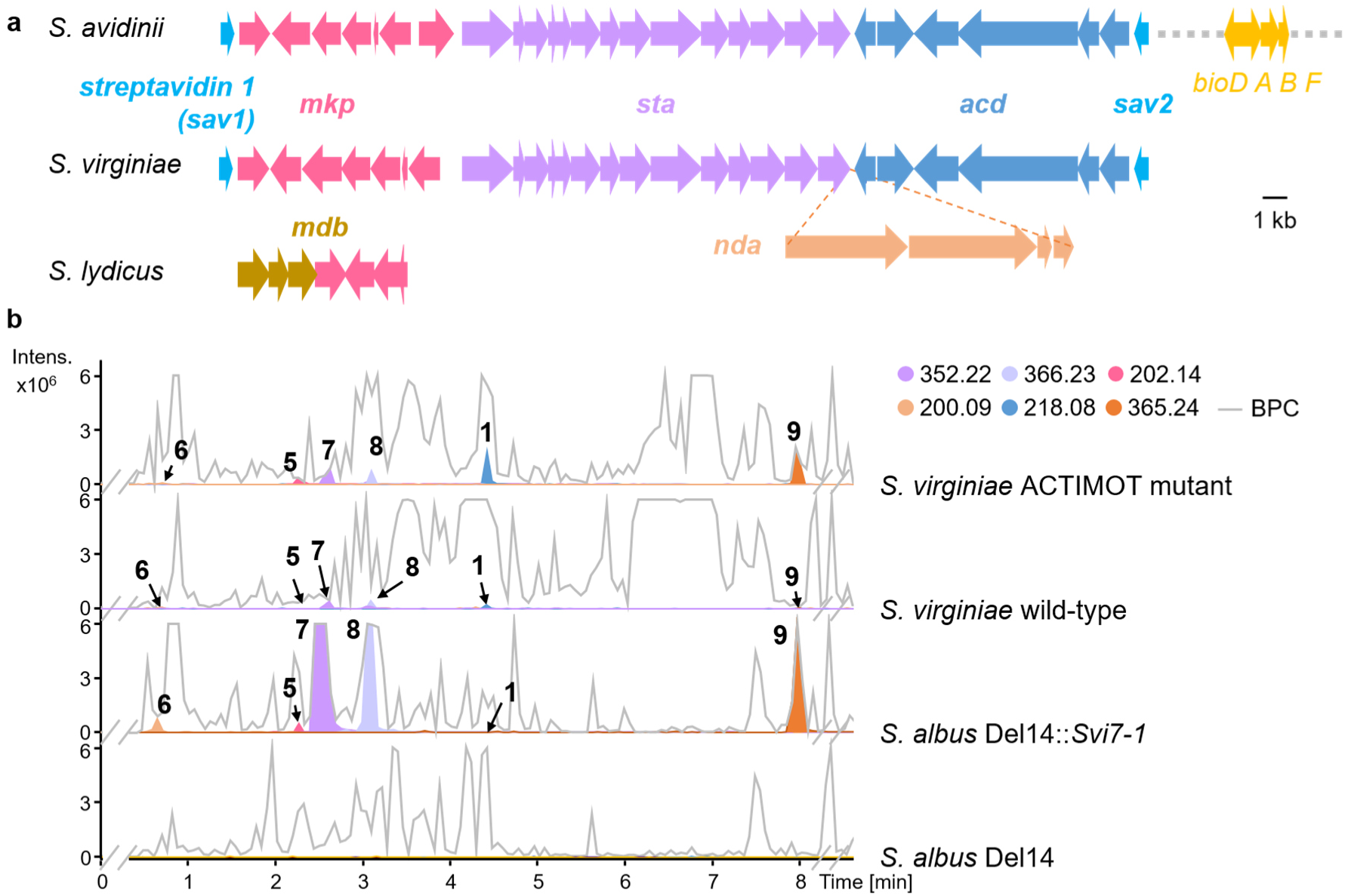
Identification and analysis of the biotin-related “BGC Nexus” and its activation in *S. virginiae* using ACTIMOT. **a**, Organization of the biotin-related “BGC Nexus” flanked by two streptavidin genes (*sav1* and *sav2*, azure). Three conserved BGCs for α-methyl-KAPA (*mkp,* magenta), stravidins^24,25^ (*sta*, light purple), and acidomycin (*acd,* light navy) are present in both *S. avidinii* DSM40526 and *S. virginiae* DSM40094. *S. virginiae* harbors one additional BGC (*nda*, light orange) responsible for a new non-proteinogenic amino acid, 2-aminonona-5,7-dienedioic acid (ANDA). In *S. lydicus* DSM40461, which lacks streptavidin genes, the *mkp* cluster is conserved together with an additional copy of the *bioADB* operon (dark gold), termed the *mdb* cluster. **b**, Base peak chromatograms (BPCs) and extracted ion chromatograms (EICs) of culture extracts from wild-type and ACTIMOT-engineered *S. virginiae* and *S. albus* Del14 strains are shown. Targeted mass-to-charge (*m/z*) values corresponding to the detected metabolites are indicated in each trace.

Closer inspection of this region revealed two additional putative BGCs flanking *sta*, all positioned between *sav1* and *sav2*, forming a dense genomic region we term the “BGC Nexus.” A comparable architecture was found in *S. virginiae* DSM40094, the acidomycin producer^9^, which contains the same core arrangement plus an additional BGC (designated *nda*, Fig. 2a, light orange). The consistent co-localization of multiple BGCs with dual streptavidin genes and the primary biotin operon suggests an evolutionarily conserved genomic organization that encodes a coordinated system of biotin-related metabolites and their associated binding proteins (Fig. 2a).

### ACTIMOT enables discovery of products from the “BGC Nexus” in *S. virginiae*

To elucidate the metabolic output of the four BGCs within the “BGC Nexus”, we employed ACTIMOT^23^ to mobilize the target DNA region *Svi7-1* from *S. virginiae*, which encompasses all BGCs, yielding the multi-copy construct pCap-*Svi7-1* (Supplementary Figs. 1-2). The resulting ACTIMOT mutant and the wild-type (WT) strain were cultivated under identical conditions, and their extracts were analyzed by ultra-performance liquid chromatography-high-resolution mass spectrometry (UPLC-HRMS) (Fig. 2b). Compared with the WT, the ACTIMOT mutant showed moderately increased production of the unknown compound **6** and known stravidins (**7** and **8**, Supplementary Figs. 3-5, 19, 29-32, Supplementary Table 6), and a pronounced accumulation of three additional metabolites (**1**, **5**, and **9**). HRMS analysis suggested that **1** corresponds to acidomycin (Supplementary Figs. 3-5), a known BioB inhibitor^19^ whose biosynthetic origin has remained elusive. These results indicated that **1**, **5**, **6**, and **9** originate from the three uncharacterized BGCs. Heterologous expression of pCap-*Svi7-1* in *S. albus* Del14 reproduced all six compounds, confirming activation of the complete “BGC Nexus” (Fig. 2b). Subsequently, our focus shifted toward identifying these products and establishing their linkage to the corresponding BGCs.

### The *mkp* BGC is responsible for α-methyl-KAPA production

Among the four BGCs uncovered in the *S. virginiae* “BGC Nexus”, the *mkp* cluster attracted particular attention due to its atypical gene composition. The *mkp* cluster comprises seven genes (*mkpA–G*) encoding enzymes related to fatty acid biosynthesis and tailoring modifications (Fig. 3a,c, Supplementary Table 4). This cluster was cloned into the p15A backbone under the *kasO*p*\** promoter, yielding p15A-*kasO*p*\*-mkp* (Fig. 3a). Heterologous expression of p15A-*kasO*p*\*-mkp* in *S. albus* Del14 led to the production of compounds **5** and **9** (Fig. 3b). NMR structural analysis identified compound **5** as α-methyl-KAPA and compound **9** as its symmetric dimer, formed spontaneously from **5** (Supplementary Figs. 3-4, 20-21, 33-42, Supplementary Tables 7-8).

**Fig. 3:**
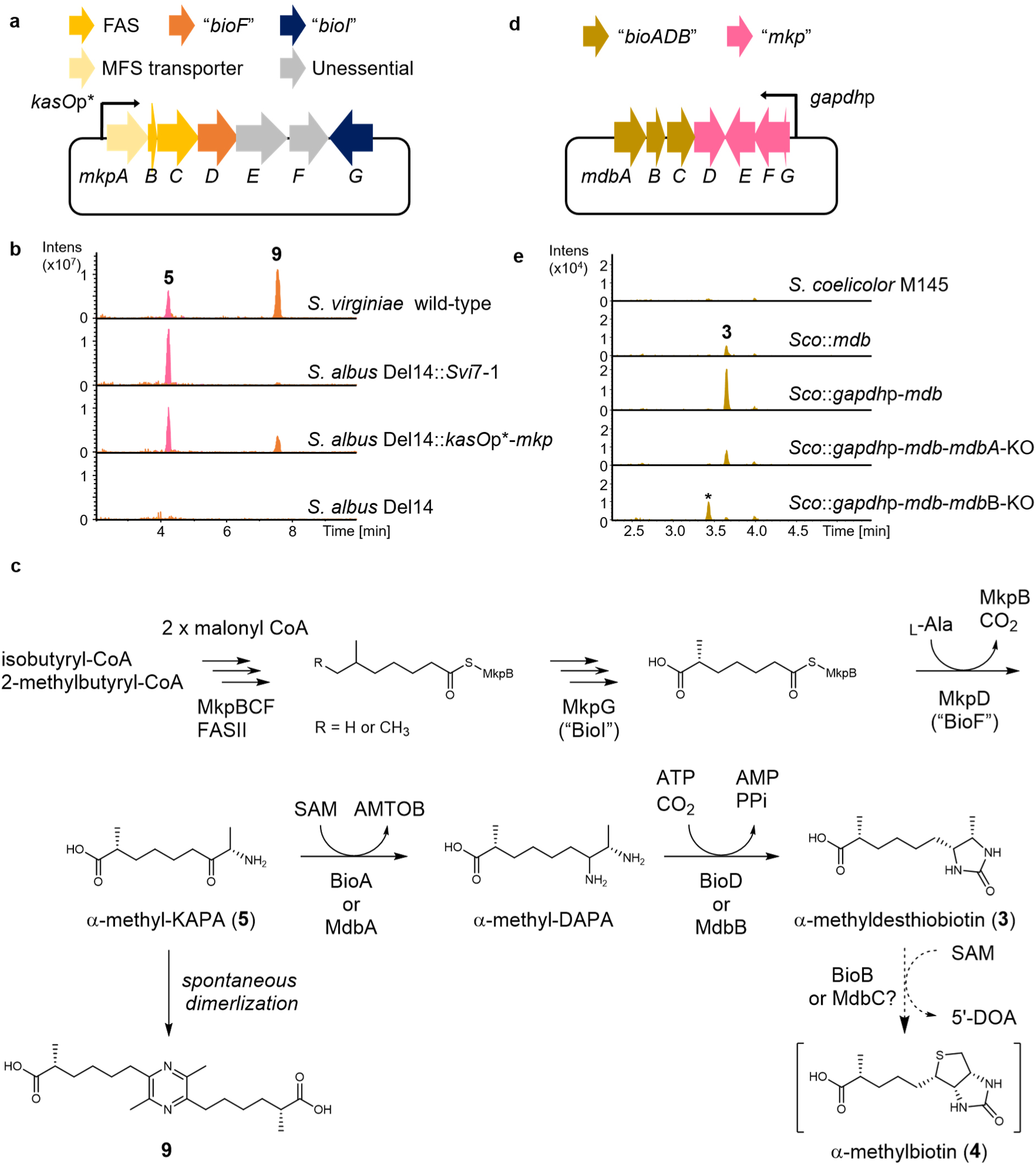
Heterologous production and biosynthetic pathways of α-methyl-KAPA and α-methyldesthiobiotin. **a**, Plasmid construct for heterologous expression of the *mkp* cluster. **b**, LC-MS analysis of the heterologous expression strains. EICs of *m/z* 202.14 and 365.24 corresponding to α-methyl-KAPA (**5**, magenta) and the oxidized dimer (**9**, orange) are shown. **c**, Proposed biosynthetic pathway for **5** and **3**. KAPA, 7-keto-8-aminopelargonic acid; DAPA, 7,8-diaminopelargonic acid; SAM, *S*-adenosyl methionine; AMTOB, *S*-adenosyl-2-oxo-4-methylthiobutyric acid; 5′-DOA, 5′-deoxyadenosine. **d**, Plasmid construct for heterologous expression of the *mdb* cluster, comprising “*bioADB*” together with the *mkp* cluster derived from *S. lydicus.* **e**, LC-MS analysis of heterologous expression and gene knockout mutants. The EICs for *m/z* 229.16 corresponding to **3** are shown in dark gold. *Sco* denotes *S. coelicolor* M145. KO indicates gene knockout. The peak marked with an asterisk represents an isobaric compound not related to α-methyldesthiobiotin.

Bioinformatic annotation indicated that MkpB and MkpC function as an acyl carrier protein (ACP) and a ketosynthase, respectively, while MkpF shares 56% sequence similarity with FabH (β-ketoacyl-ACP synthase III)^26^ (Supplementary Table 4). We propose that endogenous FabD from the fatty acid synthase (FAS) pathway transfers malonyl-CoA to MkpB, followed by condensation with isobutyryl-CoA or 2-methyl-butyryl-CoA catalyzed by MkpF, and two successive chain-elongation cycles mediated by MkpC (Fig. 3c). The resulting intermediate is predicted to undergo oxidative cleavage by MkpG, a cytochrome P450 homologous to BioI from *Bacillus subtilis*^27^, producing a 6-methyl-pimelic acid intermediate. Finally, MkpD, a BioF-like enzyme, catalyzes condensation with L-alanine and decarboxylation to **5** (Fig. 3c).

To validate this biosynthetic model, we generated individual gene deletions using Red/ET recombination^28^. Deletion of *mkpC*, *mkpD*, or *mkpG* completely abolished production of **5**, whereas disruption of *mkpE* or *mkpF* had no effect, confirming *mkpCDG* as essential genes for this pathway (Supplementary Fig. 6). The presence of **5** in the *mkpF* mutant likely results from complementation by an endogenous *fabH* homolog. Isotope-feeding experiments with labeled precursors further supported the proposed pathway: supplementation with L-valine-*d*_8_ or L-threonine-^13^C_4_,^15^N resulted in +4 Da and +2 Da mass shifts in **5** (Supplementary Fig. 7), indicating that both isobutyryl-CoA and 2-methyl-butyryl-CoA can serve as starter units in α-methyl-KAPA biosynthesis.

### α-methyl-KAPA serves as precursor for α-methyldesthiobiotin

To investigate the potential target of **5**, we employed an *in vitro* BioA-BioD coupling assay, which unexpectedly led to the formation of α-methyldesthiobiotin (**3**) (Supplementary Fig. 8). This suggested that **5** could enter the biotin biosynthetic pathway as a substrate, leading toward **3** and **4**. However, only **3**, but not **4**, was detected in extracts of both *S. virginiae* WT and ACTIMOT mutant strains (Supplementary Fig. 9), as well as in *E. coli* cultures supplemented with **5** (Supplementary Fig. 10).

Previous reports described the formation of **3** and trace detection of **4** in *S. lydicus*^17,18^. However, **4** has never been isolated as a natural product and is only available as a synthetic mixture of diastereomers^17,18^. We isolated **3** from *S. lydicus* and demonstrated it indeed is α-methyldesthiobiotin (Supplementary Figs. 3-4, 22, 43-47, Supplementary Table 9). Our bioinformatic analysis revealed that *S. lydicus* DSM40461 contains the *mkp* cluster adjacent to an additional *bioADB* operon homologous to the primary biotin operon (Figs. 2a and 3d, Supplementary 4). To examine this relationship, we heterologously expressed the “*bioADB*”*–mkp* cassette from *S. lydicus* (termed *mdb*) in *S. coelicolor* M145 resulting in the mutant *Sco*::*mdb*. UPLC–HRMS analysis showed robust production of **3** but not **4** (Fig. 3e). Overexpression of *mkp* under the *gapdh*p promoter enhanced the yield of **3**, whereas deletion of *mdbA* (“*bioA*”) or *mdbB* (“*bioD*”) decreased its production (Fig. 3e), confirming their roles in producing **3**, although the pathway can be partially complemented by primary *bioAD*.

Finally, *in vitro* reconstitution with purified MdbA and MdbB converted **5** to **3** (Supplementary Fig. 11). Together, these findings demonstrate that α-methyl-KAPA (**5**) functions as a substrate within the biotin biosynthetic pathway, leading to the formation of α-methyldesthiobiotin (**3**) (Fig. 3c).

### The optional BGC *nda* directs biosynthesis of 2-aminonona-5,7-dienedioic acid (ANDA) and its dipeptide derivative

Following the characterization of the *mkp* pathway and its metabolic link to the biotin biosynthetic route, we turned our attention to *nda*, a unique BGC present in *S. virginiae* but absent in *S. avidinii*. Located between *acd* and *sta* within the biotin-related “BGC Nexus,” *nda* encodes a compact assembly of three biosynthetic enzymes and a transporter protein, including a polyketide synthase (PKS) and a nonribosomal peptide synthetase (NRPS) (Figs. 2a and 4a, Supplementary Table 4).

**Fig. 4:**
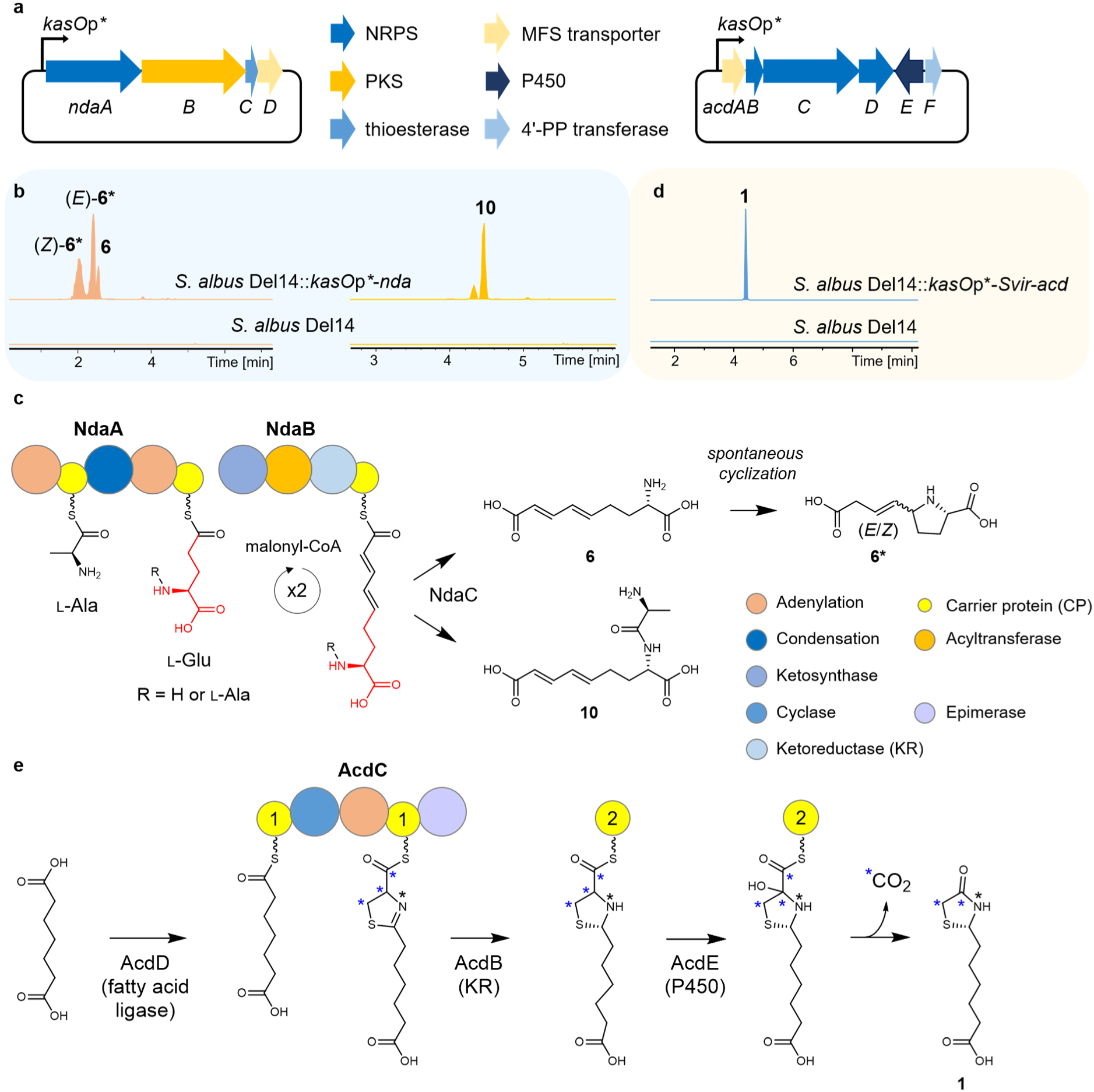
Heterologous production of ANDA and acidomycin and their proposed biosynthetic pathways. **a**, Plasmid constructs for heterologous expression of the *nda* (left) and *acd* (right) clusters. **b**, Heterologous production of ANDA (**6**), its isomers (**6\***), and the alanine dipeptide derivative Ala-ANDA (**10**). EICs of *m/z* 200.09 for **6** and **6\*** (orange) and 271.13 for **10** (blue) are shown. **c**, Proposed biosynthetic pathway for **6** and **10**. The structural moiety derived from glutamic acid is highlighted in red. Each circle represents an enzymatic domain, with annotations color-coded as indicated in the legend (right). **d**, Heterologous production of acidomycin (**1**). The EIC of *m/z* 218.09 is shown in light blue. **e**, Proposed biosynthetic pathway of acidomycin. Carbon atoms derived from cysteine are marked with asterisks (**\***) in blue.

To investigate its products, *nda* was cloned under the constitutive *kasO*p*\** promoter and heterologously expressed in *S. albus* Del14 (Fig. 4a). LC–MS analysis revealed a major compound (**6**, *m/z* 200.09) and an additional derivative (**10**, *m/z* 271.13) (Fig. 4b). Purification and structural analysis identified **6** as 2(*S*)-aminonona-5,7-dienedioic acid (ANDA) and **10** as its alanine dipeptide conjugate, Ala-ANDA (Supplementary Figs. 3-4, 23-24, 48-56, Supplementary Tables 10-11). Additionally, earlier-eluting peaks with the same *m/z* (denoted **6\***) were shown by NMR to be cyclized isomeric derivatives of **6** that form spontaneously (Fig. 4b,c and Supplementary Figs. 3-4, 25-26, 57-66, Supplementary Tables 12-13). Finally, the absolute configurations at C-2 and C-2’ in **10** were determined as 2*S*,2’*S* through ozonolysis (Supplementary Fig. 27) followed by Marfey’s method (Supplementary Fig. 12).

Stable-isotope feeding with L-glutamic acid-^13^C_5_ produced a +5 Da mass shift in **10** (Supplementary Fig. 13), demonstrating that glutamic acid constitutes the skeleton of **6** and **10**. Based on these findings, we propose that NdaA activates L-alanine and L-glutamic acid via its NRPS adenylation domains, followed by two rounds of chain extension by the PKS NdaB and release by the thioesterase NdaC, yielding **6** or **10** (Fig. 4c).

### Uncovering the biosynthetic origin for acidomycin

The remaining *acd* BGC was hypothesized to direct the biosynthesis of **1**, also detected in *S. avidinii* (Fig. 2a, Supplementary Table 4). To verify this, *acd* (six genes) from both *S. virginiae* and *S. avidinii* were cloned under the *kasO*p*\** promoter and expressed in *S. albus* Del14, yielding strains *S. albus* Del14::*kasO*p*\*-Svir-acd* and *S. albus* Del14::*kasO*p*\*-Savi-acd* (Fig. 4a). UPLC-HRMS analysis of both transformants confirmed production of **1** (Fig. 4d and Supplementary Fig. 14). NMR and optical rotation analyses established the structure of **1** and, for the first time, unambiguously identified the natural compound as (*S*)-(−)-acidomycin (Supplementary Figs. 3-4, 28, 67-71, Supplementary Table 14).

We proposed that pimelic acid serves as the starter unit in the biosynthesis of **1**. AcdD, annotated as AMP ligase, activates pimelic acid through adenylation and subsequently loads it onto the first carrier protein domain (CP1) of AcdC (Fig. 4e). The adenylation domain of AcdC adenylates and attaches cysteine to the CP2. This is followed by condensation and cyclization by the cyclization domain, resulting in the thiazoline ring formation. AcdB, annotated as ketoreductase, reduces the double bond to yield a thiazolidine moiety, while AcdE, a P450 hydroxylase, potentially introduces a hydroxyl group at the α-carbon. The product is proposed to be released from AcdC through oxidative decarboxylation, possibly mediated by a non-canonical activity of AcdC or AcdE, yielding **1** (Fig. 4e). This proposed pathway gained further support from an isotope-feeding experiment, in which a mass shift of +3 Da for acidomycin was observed upon supplementation of L-cysteine-^13^C_3_,^15^N into the culture of *S. avidinii-kasO*p*\*-aviA* (Supplementary Fig. 15).

### Anti-bacterial activities of products from the biotin-related “BGC Nexus”

While **5** has been identified as an inhibitor of leukotriene-A4 hydrolase^20^, its structural similarity to KAPA suggested that it could act as a competitive inhibitor of BioA and possibly other enzymes in the biotin biosynthetic pathway. To test this hypothesis, we conducted bioactivity assays with *E. coli* BW25113 and its Δ*bioF* mutant^29^ in M9 minimal media in the absence of biotin. As the biotin biosynthetic pathway is strictly regulated by the repressor BirA which functions sensitively depending on the concentration of biotin^7,30^, cells were cultured for four hours before **5** was added to the culture to allow for biotin consumption^31^. After an additional 19 hours of incubation, **5** exhibited a minimum inhibitory concentration (MIC) of BW25113 wild-type at 128 μM and of strain Δ*bioF* at 2.1 μM (Fig. 5b). In addition, 1 μM of DAPA rescued the growth of Δ*bioF* in the presence of **5**, while 0.1 μM of KAPA did not. The latter concentration was sufficient to rescue the growth of Δ*bioF* in the absence of **5**, proving **5** as a new BioA inhibitor (Supplementary Fig. 16). The antibacterial activities of **1**, **3**, **4**, **6**, **6\***, and **10** were also tested in a similar manner. The antibacterial activities for **1** and **3** were confirmed as reported previously, while inhibitory activities of **4**, **6**, **6\***, and **10** were not found for *E. coli* (Fig. 5b).

**Fig. 5:**
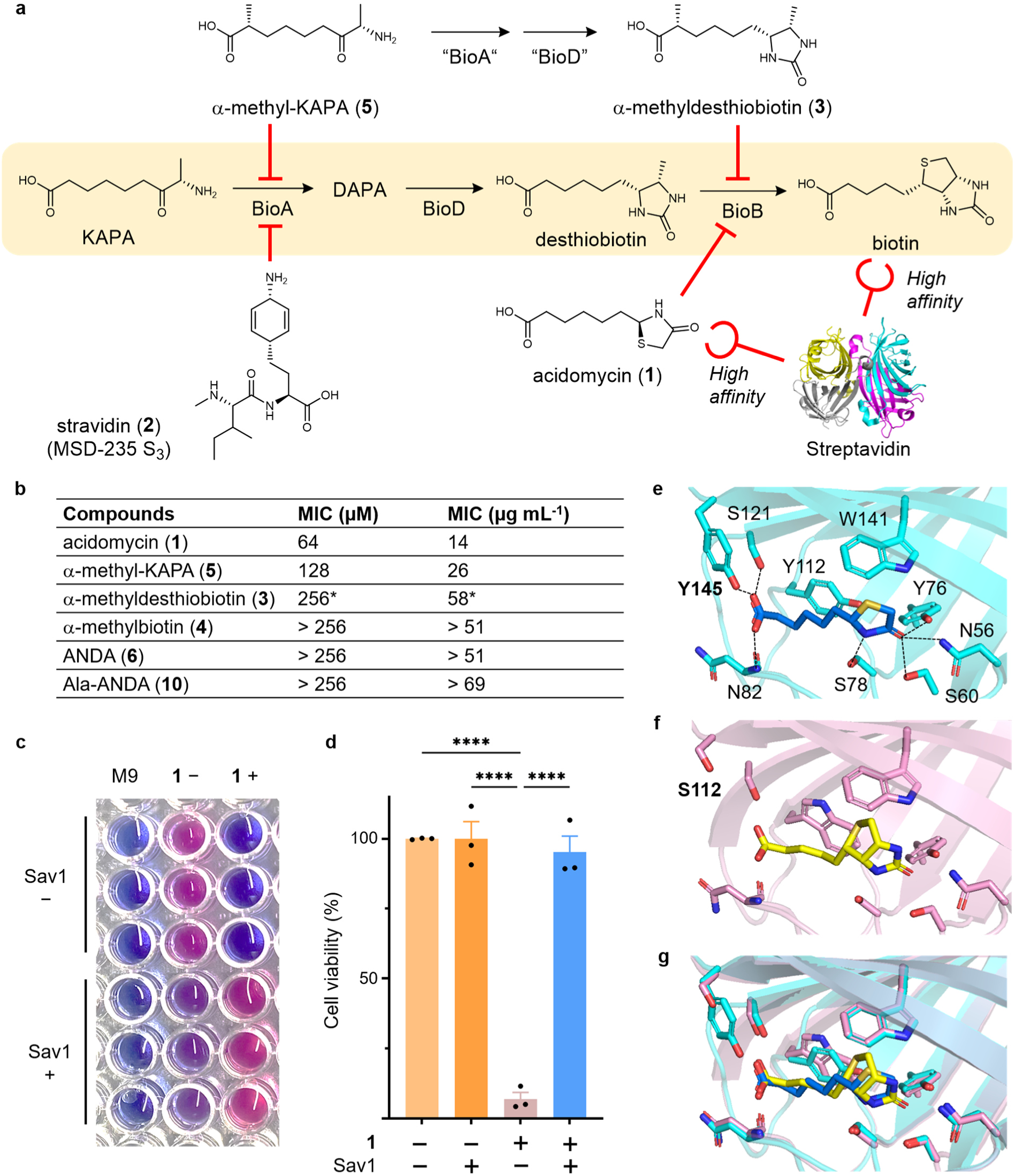
Antibacterial activities of biotin-related antibiotics and the interaction with streptavidin. **a**, The biotin-related compounds produced by the “BGC Nexus” and the interaction between streptavidin and biotin or acidomycin (**1**). The biotin biosynthetic pathway is highlighted in yellow. **b**, Antibacterial activities against *E. coli* BW25113 measured in M9 media. An asterisk denotes the concentration showing a 66% growth inhibition instead of the standard MIC. **c**, Cell viability visualization using resazurine. Wells in each lane contain M9 media (M9), *E. coli* BW25113 without (**1**−) or with 64 µM acidomycin (**1**+) at the final concentration. The top three rows contain PBS buffer as a control without adding streptavidin 1 (Sav1 −), and the bottom three rows contain 131 µM streptavidin 1 (Sav1 +) at the final concentration in PBS buffer. **d**, Statistical analysis of the effect of Sav1 on the antibacterial activity of **1**. Three biological replicates were performed, and error bars represent the means ± SEM. Statistical significance was determined by one-way ANOVA with multiple comparisons. ∗∗∗∗*p* < 0.0001. **e**, Crystal structure of the streptavidin 2 (Sav2)–acidomycin (**1**) complex (PDB ID: 9XIK), with Sav2 shown in cyan and **1** in marine. **f**, Reported crystal structure of the Sav1–biotin complex (PDB ID: 1STP)^22^, with Sav1 in pink and biotin in yellow. Hydrogen bonds are indicated by black dashed lines. **g**, Superimposed structures of Sav2-**1** and Sav1-biotin.

To further examine the inhibitory activity of **5** *in vitro*, we performed a coupled assay of recombinant *E. coli* BioA and BioD using KAPA as the substrate. LC-MS analysis of the reaction mixture showed that the production of desthiobiotin was not significantly changed, however, α-methyldesthiobiotin (**3**) was produced by adding **5** to the reaction mixture (Supplementary Fig. 8). These results indicated that **5** can be metabolized and function as a competitive inhibitor for the last steps of biotin biosynthesis. We next analyzed the metabolites of *Streptomyces* strains harboring the *mkp* cluster and from *E. coli* cultures incubated with **5**. In all cases, LC–MS consistently detected production of **3** but not **4** (Fig. 3e, Supplementary Figs. 9-10), demonstrating that **5** is indeed converted endogenously into **3**. The absence of **4** suggests that MdbC (“BioB”) in *S. lydicus* may function as a self-resistance protein and is only weakly active in this transformation under certain conditions, given that the only reported observation of natural **4** was only in trace amount^17,18^, which needs further investigation.

### Interplay of streptavidin and biotin-related antimetabolites

In addition to producing diverse biotin-related antimetabolites, another striking feature of this “BGC Nexus” is the presence of two highly conserved *streptavidin* genes, *sav1* and *sav2*. In *S. avidinii*, *sav1* encodes the canonical, structurally characterized streptavidin (PDB ID: 1STP)^22^, while *sav2* encodes a new streptavidin sharing 75% sequence identity with Sav1. Considering that *sav2* is located adjacent to the BGC for acidomycin (**1**) which shows partial structure similarity to biotin, we hypothesized that streptavidins might directly interact with **1**.

To test this, we evaluated the antibacterial activity of **1** in the presence of Sav1. Remarkably, the addition of Sav1 almost completely abolished the activity of **1**, suggesting high-affinity binding analogous to the canonical biotin–streptavidin interaction (Fig. 5c,d). To elucidate the structural basis of this interaction, we crystallized Sav2 in complex with **1** and resolved its structure at 1.8 Å resolution in a *I*4_1_22 space group (Supplementary Table 5). Interestingly, although Sav2 proteins were stable tetramers in solution, the asymmetric unit in the crystal lattice contained only one Sav2 molecule, closely resembling the Sav1 (with their RMSD of 0.9 Å) (Supplementary Fig. 17). Distinct electron density corresponding to **1** was observed at the canonical biotin-binding pocket (Supplementary Figs. 17 and 18a), confirming tight binding of **1**. The stereochemistry of the 2-pyrrolidinone moiety in **1** was determined to be *S,* which agreed with our structural analysis.

In the complex, the ɷ-carboxy group of **1** forms hydrogen bonds with Asn82, Ser121, and Tyr145, while its 2-pyrrolidinone amine and carbonyl groups are stabilized by Ser78 and Asn56/Ser60/Tyr76, respectively (Fig. 5e, Supplementary Fig. 18b). The hydrophobic residues Tyr112 and Trp141 shape the binding pocket. Interestingly, Tyr145 in the Sav2-**1** complex is replaced by the structurally smaller residue Ser112 in the Sav1-biotin complex (Fig. 5f,g). This amino acid substitution may subtly modulate binding affinity and facilitate competitive biotin sequestration in the environment, as **1** does not have a spatial ring structure such as the sulfur-containing tetrahydrothiophene ring in biotin.

## Discussion

The biotin biosynthetic pathway represents an attractive antibacterial target due to its essentiality in bacteria and absence in humans^1–3^. In this study, we uncovered a conserved genomic architecture, a biotin-related “BGC Nexus”, comprising three to four BGCs flanked by two streptavidin genes (Fig. 2a). This genomic configuration, recurrently found across producers of known biotin antimetabolites including *S. avidinii* (stravidins and amiclenomycin), *S. virginiae* (acidomycin), and *S. lydicus* (α-methylated analogues), highlights an evolutionary strategy that links biotin-binding proteins with multiple biosynthetic pathways targeting biotin metabolism (Fig. 5a). Activation of this region in *S. virginiae* using ACTIMOT^23^ revealed a chemically and functionally diverse repertoire of biotin antimetabolites, revealing that the streptavidin-associated genomic regions encode a previously hidden layer of metabolic complexity (Fig. 2b).

Among these metabolites, stravidins, encoded by the *sta* cluster, exemplify a rare form of cooperative antibiotic system involving both small molecules and proteins. Historical studies demonstrated that streptavidin potentiates the antibacterial activity of stravidins^14–16^, which inhibit BioA through covalent modification of its pyridoxal phosphate cofactor^32–35^. This synergy likely reflects a dual mechanism: streptavidin depletes extracellular biotin, sensitizing competitors to inhibition of the biotin pathway, while stravidins directly block a key enzymatic step (Figs. 5a and 6).

**Fig. 6:**
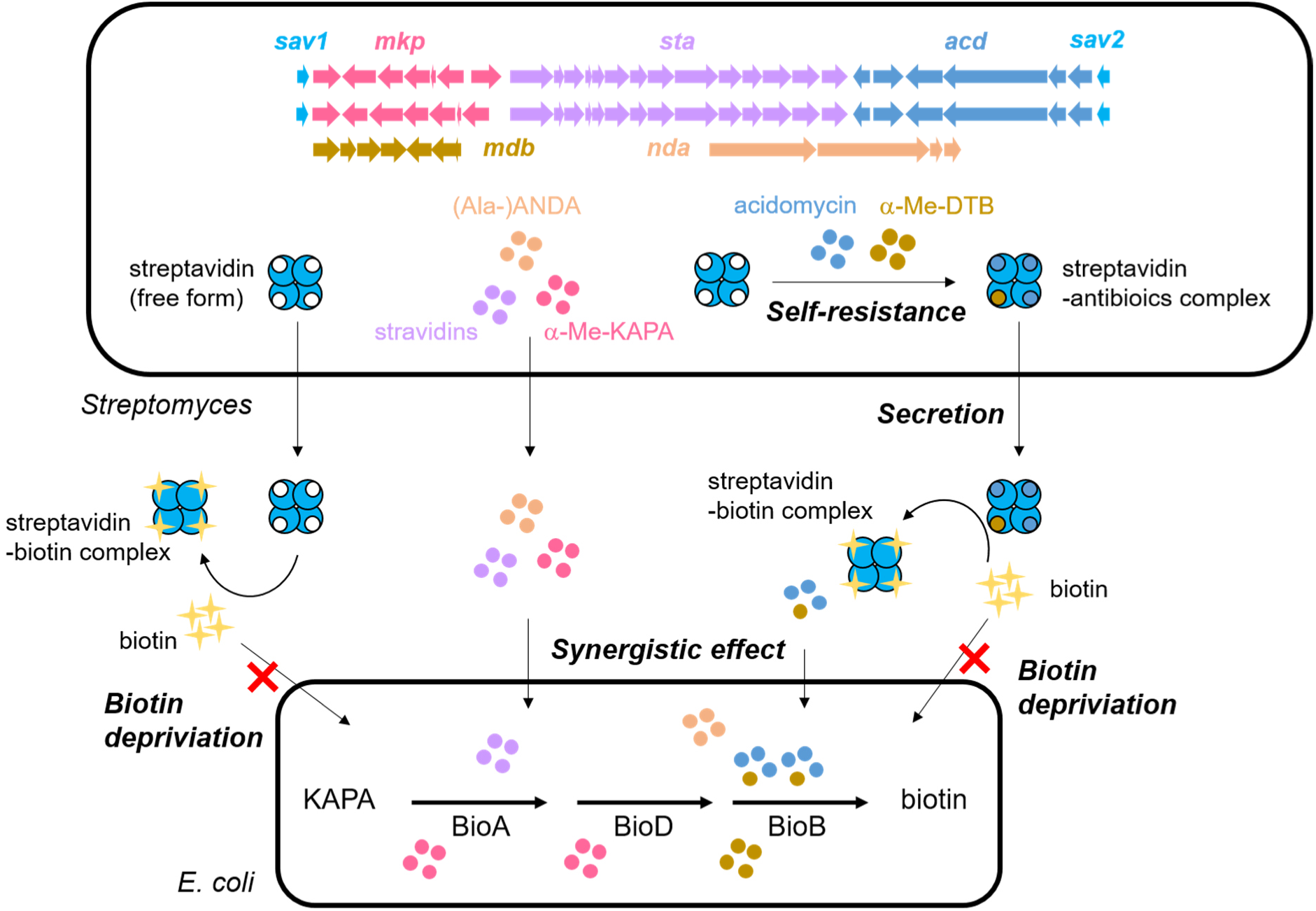
Proposed model for the functions of streptavidin in the biotin-related antibiotic network. Streptavidins (azure) capture acidomycin (light navy) and possibly α-methyldesthiobiotin (α-Me-DTB, dark gold), providing the producing organism with self-resistance against its own biotin-pathway antimetabolites. The streptavidin–antimetabolite complex is secreted into the environment, where streptavidin acts as a biotin scavenger, triggering compensatory biotin biosynthesis in neighboring bacteria. The resulting biotin deficiency sensitizes competing cells to biotin-pathway inhibitors such as α-methyl-KAPA and stravidins, culminating in bacterial growth inhibition and ecological dominance of the producer.

We further identified α-methyl-KAPA (**5**), produced by the *mkp* cluster, as a previously unrecognized BioA inhibitor and a prodrug-like precursor that enters the biotin biosynthetic pathway, where it is enzymatically converted into α-methyldesthiobiotin (**3**). This represents the first experimental evidence that α-methyl-KAPA directly targets BioA, revealing its dual role as both an enzyme-directed antimetabolite and a biosynthetic intermediate. Although α-methylbiotin (**4**) has historically been proposed as a related metabolite, we consistently observed only **3** in both *in vitro* and *in vivo* systems, aligning with earlier observations that **4** appears only in trace amounts in *S. lydicus* fermentation and has been characterized only via total synthesis as a diastereomeric mixture^17,18^. Given that **3** is a known BioB inhibitor^19^, these data support a model in which **3**, rather than **4**, represents the physiologically active species. This metabolic conversion, from α-methyl-KAPA to α-methyldesthiobiotin, demonstrates a remarkable interplay between primary and secondary metabolism, where intermediates of core biosynthetic pathways are repurposed into potent natural antimetabolites.

The *nda* cluster, unique to *S. virginiae*, encodes the biosynthesis of the non-proteinogenic amino acid ANDA and its alanine dipeptide derivative. Although their precise roles and molecular targets remain unknown (Fig. 5b), these compounds expand the structural diversity of biotin-pathway-related natural products, consistent with their localization within the biotin-associated “BGC Nexus.” In parallel to our work, a recent preprint by Brown and colleagues described related genomic loci in other *Streptomyces* species, identifying comparable classes of biotin-related metabolites, including ANDA-like compounds, and proposed BioD as a potential target, although this hypothesis still awaits direct experimental validation^36^. Together, these independent findings collectively reinforce the concept that ANDA and its derivatives may function as biotin antimetabolites and underscore the broad phylogenetic conservation of this metabolic strategy.

The *acd* cluster, responsible for acidomycin biosynthesis, provides a striking example of how nature repurposes core metabolic scaffolds. A recent report proposed that acidomycin (**1**, also referred to as mycobacidin) arises from the activity of BioADB enzymes within the primary biotin operon of *S. virginiae*^37^. However, that mechanism likely reflects formation of **1** from the chemically synthesized 7-oxoheptanoate precursor, potentially via side reactions catalyzed by BioADB. In contrast, the clear and reproducible production of **1** through our heterologous expression of *acd* from multiple strains provides definitive evidence for its most likely genuine biosynthetic origin.

It was reported that acidomycin inhibits BioB under biotin-depleted conditions^38^. However, unlike stravidins^15^, our results showed that the activity of acidomycin is attenuated by streptavidin. Crystallographic analysis demonstrated that acidomycin binds streptavidin at the canonical biotin-binding pocket, adopting a near-identical conformation to biotin (Fig. 5e-g). These findings suggest that streptavidin is a multifunctional protein. Beyond serving as a biotin scavenger, it can bind and neutralize acidomycin, thereby conferring self-protection to the producer. Based on these observations, we propose a working model in which the streptavidin–acidomycin complex may be exported from the cell, where environmental biotin could displace acidomycin, releasing the active compound to inhibit BioB in competing bacteria. Concurrently, freed streptavidin resumes its role as a high-affinity biotin binder, potentially exacerbating local biotin limitation and sensitizing neighboring cells to co-produced inhibitors (Fig. 6). Although this model aligns with the synergistic behavior of biotin-pathway antimetabolites observed under defined biotin-depleted conditions^36^, direct evidence for streptavidin-mediated regulation or ecological coordination remains to be established.

Collectively, our findings suggest that *Streptomyces* species have evolved a genomic and biochemical framework that links biosynthetic enzymes with binding proteins to manage biotin-related metabolites and mitigate self-toxicity. The streptavidin-flanked “BGC Nexus” provides a structural and functional context for such interactions, connecting protein binding with metabolic diversification and potentially facilitating the coexistence of biotin metabolism and antimetabolite production. More broadly, this work highlights the power of ACTIMOT to uncover hidden biosynthetic potential and reveal the complexity of metabolite–protein cooperation in natural product systems. The discovery of the streptavidin-anchored “BGC Nexus” extends the paradigm of microbial secondary metabolism beyond discrete gene clusters to higher-order genomic architectures that integrate chemical and protein-mediated defense.

## Methods

### General materials and procedures of microbiology, molecular biology, and bioinformatics

Bacterial strains and plasmids used in this study are listed in Supplementary Tables 1 and 2, and primers are provided in Supplementary Table 3. DNA polymerases, restriction enzymes, T4 DNA ligase, plasmid extraction kits, and other routine molecular biology reagents were obtained from Thermo Scientific. Gibson assemblies were carried out using the NEBuilder HiFi DNA Assembly Kit (New England Biolabs Inc.). SuperCos 1 Cosmid Vector Kit was purchased from Agilent Technologies. Standard molecular cloning procedures—including PCR, restriction digestion, ligation, Gibson assembly, and plasmid extraction—were performed according to manufacturers’ instructions. Plasmid modifications by Red/ET recombination followed established protocols^39,40^. α-Methyl-KAPA and α-methylbiotin were purchased from Enamine Ltd. (Ukraine) and Toronto Research Chemicals (LGC, Canada), respectively.

Genomic DNA from *Streptomyces* strains was extracted using a cetyltrimethylammonium bromide (CTAB)-based method^41^. Actinobacterial strains were maintained at 30 °C on ISP4 or MS agar plates, while *E. coli* strains were cultured in Luria–Bertani (LB) medium. For preparing the ISP4 medium, two solutions are required. Solution I: make a paste of 10.0 g starch with a small amount of cold deionized distilled water (ddH_2_O) and fill the volume up to 500 mL. Solution II: 2 g CaCO_3_, 1 g K_2_HPO_4_ (anhydrous), 1 g MgSO_4_ x 7 H_2_O, NaCl 1g, 2 g (NH_4_)_2_SO_4_, 500 mL ddH_2_O, 1 mL trace salt solution (0.1 g FeSO_4_ x 7 H_2_O, 0.1 g MnCl_2_ x 4 H_2_O, 0.1 g ZnSO_4_ x 7 H_2_O, 100 mL ddH_2_O). pH was not adjusted if between 7.0 and 7.4. Mix solutions I and II. MS agar contained 20 g mannitol, 20 g soya flour, and 20 g agar per liter of tap water. Each liter of LB medium comprised 10 g tryptone, 5 g yeast extract, and 5 g NaCl (pH 7.2). *In silico* analyses, including primer design, BGC annotation, and short-read mapping, were performed using Geneious Prime® 2024.0.7 (Biomatters Ltd., New Zealand).

### Mobilization, relocation, multiplication, and recovery of *Svi7-1* in *S. virginiae*

For mobilization and multiplication of the target DNA region (TDR) *Svi7-1* from *S. virginiae*, a single working plasmid, pRelCap-SVI7-1-dsp, was constructed following the standard ACTIMOT protocol^23^. In brief, two basic plasmids, pRel-SVI7-1-dsp and pCap-SG5-Apr-SVI7-1-LR, were first generated and subsequently fused to yield pRelCap-SVI7-1-dsp.

To construct pRel-SVI7-1-dsp, primer pairs of two spacers (*Svi7-1*-sp1 and *Svi7-1*-sp2) were annealed and ligated to an *Nco*I + *Xba*I pre-treated pRel or pHelp vectors to produce pRel-SVI7-1-sp1 or pHelp-SVI7-1-sp2, respectively. The pRel-SVI7-1-sp1 was then digested overnight with *Nhe*I, *Bcu*I, and *Stu*I, and the resulting 5.5-kb and 4.4-kb fragments were purified from agarose gel. In parallel, *Svi7-1*-DelL, *Svi7-1*-DelR, and the second sgRNA cassette were amplified from the genomic DNA and pHelp-SVI7-1-sp2, respectively. The plasmid pRel-SVI7-1-dsp construct was then constructed via Gibson assembly using these five fragments obtained above. For pCap-SG5-Apr-SVI7-1-LR, the amplified *Svi7*-*1*-CapL and *Svi7-1*-CapR fragments were assembled with the *Mss*I pre-treated pCap-SG5-Apr through Gibson assembly. Finally, both pRel-SVI7-1-dsp and pCap-SG5-Apr-SVI7-1-LR were treated with *Mss*I, and the purified fragments were combined via Gibson assembly to generate pRelCap-SVI7-1-dsp.

The working plasmid pRelCap-SVI7-1-dsp was introduced into *S. virginiae* via conjugation as described below. The resulting exconjugants were picked and cultivated in TSB for three days. Total plasmids were isolated from the ACTIMOT mutants using a modified alkaline lysis method previously described^23^. The isolated plasmid samples were subsequently electroporated into *E. coli* DH10B. Single *E. coli* transformants were cultured overnight in LB liquid supplemented with apramycin and cultivated for plasmid amplification and extraction. The presence of pCap-*Svi7-1* was verified by *Eco72*I restriction digestion (Supplementary Fig. 1) and confirmed through short-read sequencing (Supplementary Fig. 2).

### *S. avidinii* DSM 40526 cosmid library construction and screening

A genomic cosmid library of *S. avidinii* DSM 40526 was constructed using the SuperCos 1 vector. High-molecular-weight DNA was partially digested with *Sau3A*I, and ∼40 kb fragments were gel-purified and ligated into the prepared vector according to the SuperCos 1 kit protocol. Recombinant cosmids were transfected and arrayed in 384-well plates. Target gene clusters were identified by two-dimensional (2D) PCR screening (primers are listed in Supplementary Table 3) as described previously^42^ Positive hits were validated by end sequencing and diagnostic restriction analysis.

### Heterologous expression of BGC *mkp*, *acd* and *nda* from *S. virginiae*

For heterologous expression of the BGCs *mkp* and *nda* from *S. virginiae*, plasmids with p15A-int backbone harboring the relevant BGCs were constructed. The p15A-int backbone was amplified using primers p15A-pep-f/r. The *in silico* annotated BGCs, including the upstream constitutive promoter *kasO*p* introduced by Red/ET, were divided into 4-5 kb fragments and amplified with relevant primer pairs (Supplementary Table 3). The backbone and the fragments for each BGC were assembled using Gibson assembly to yield the corresponding plasmid for heterologous expression.

For expression of the *acd* BGC, *acdA*–*acdE* were divided into two fragments and amplified with the appropriate primers (Supplementary Table 3). These fragments were assembled with the p15A-int-kasO-sfp-amp backbone using the Gibson assembly, after which *bioF* and its native promoter were inserted at the Mph1103I site. The resulting plasmid was introduced into *S. albus* Del14 via tri-parental conjugation as described below.

### Heterologous expression of BGC *acd* from *S. avidinii*

The φC31-integrase–apramycin resistant gene (Apr^R^) cassette was inserted into SuperCos-A24, which contains BGC *acd*, by Red/ET recombineering to generate SuperCos-A24-int. A BspT1– chloramphenicol-resistant gene (Chl^R^)–BspT1 cassette was then integrated into SuperCos-A24-int to produce the minimized SuperCos-A24-int-mini, retaining *acdA–acdF* and one downstream redundant gene. This construct was further digested with BspT1 to remove the Chl^R^ cassette, and the resulting fragment was assembled with the *kasO*p* promoter fragment using Gibson assembly to yield SuperCos-*kasO*p*-*Savi-acd*. The resulting plasmid was introduced into *S. albus* Del14 via tri-parental conjugation to generate *S. albus* Del14::*kasO*p\**-Savi-acd*.

### Heterologous expression of BGC *mdb* from *S. lydicus*

To clone the *mdb* BGC from *S. lydicus*, the BGC was divided into three fragments (each 1-4 kb in size) and amplified using primers listed in Supplementary Table **3**. The amplified fragments were subsequently assembled into the vector p15A-in via Gibson assembly to yield p15A-*mdb*. Subsequently, a Chl^R^-*gapdh*p cassette was introduced into the upstream of *mdbG* by Red/ET recombination to place the constitutive promoter *gapdh*p, generating p15A-*gapdh*p-*mdb*.

For gene knockout, *mdbA* or *mdbB* was replaced with the ampicillin-resistant gene (Amp^R^) using Red/ET, resulting in p15A-*gapdh*p-*mdb*-*mdbA*KO and p15A-*gapdh*p-*mdb*-*mdbB*KO, respectively. All the constructs were introduced into *S. coelicolor* M145 for heterologous expression via bi-parental conjugation, as described below.

### Production optimization of acidomycin in *S. avidinii*

To optimize the production of acidomycin in *S. avidinii*, the native promoter of *acdA* was replaced with the strong promoter *kasO*p* through pQS-gusA-based CRISPR-Cas9 system^43,44^. The working plasmid pQS-gusA-acdA-kasOp was constructed by inserting the annealed guide duplex into *Nco*I/*Xba*I-digested pQS-gusA, followed by Gibson assembly of a pair of 2-kb homologous arms and the *kasO*p* promoter cassette using Gibson assembly. The resulting construct pQS-gusA-*acdA*-*kasO*p* was introduced into *S. avidinii* WT to generate *S. avidinii-kasOp*-acdA*, which was subsequently used for acidomycin purification and isotope-labeling experiment.

### Targeted gene deletion in the *mkp* BGC

A 26.9 kb genomic fragment from *S. virginiae*, containing the complete *mkp* BGC, was cloned into the p15A-cm-tetR-tetO-hyg-ccdB backbone using a direct-cloning approach^28^. Target genes within the cluster were replaced by an apramycin-resistance cassette via Red/ET recombination^39,40^ to generate the corresponding knockout plasmids. These plasmids were introduced into *S. virginiae*, and *in situ* gene deletions were obtained through double-crossover homologous recombination.

### Conjugations

Exogenous DNA was introduced into *Streptomyces* strains via adapted bi- or tri-parental conjugation protocols^41^. For bi-parental conjugation, *E. coli* ET12567/pUZ8002 served as the donor, whereas tri-parental conjugation employed methylation-deficient *E. coli* SCS110 as the donor and *E. coli* HB101/pRK2013 as the helper strain. Fresh *Streptomyces* spores, harvested after 3–5 days of growth, were washed twice with 2× YT medium (16 g L^-1^ tryptone, 10 g L^-1^ yeast extract, 5 g L^-1^ NaCl) and heat-shocked at 50 °C for 10 min before mating.

Donor and helper *E. coli* strains were grown overnight, diluted 1:100 in LB with appropriate antibiotics, and harvested at OD₆₀₀ = 0.4–0.6. After two washes with LB, *E. coli* cultures were mixed with *Streptomyces* spores.

Mating mixtures were plated on MS agar or ISP4 agar, incubated at 30 °C for 16–20 h, and overlaid with 1 mL sterile H₂O containing nalidixic acid (1000 µg) and apramycin (1000 µg). Plates were further incubated at 30 °C until exconjugants appeared (3–5 days).

### Fermentation of the actinobacterial strains

For small-scale fermentation, fresh actinobacterial spores were harvested from agar plates, and inoculated into 50 mL TSB medium and incubated at 30 °C with agitation (180 rpm) for 3–5 days. The seed culture was then inoculated (5%, v/v) in M2 medium (10 g L^-1^ soluble starch, 10 g L^-1^ glucose, 10 g L^-1^ glycerol, 2.5 g L^-1^ corn steep liquid, 5 g L^-1^ peptone, 2 g L^-1^ yeast extract, 1 g L^-1^ NaCl and 3 g L^-1^ CaCO_3_, pH 7.0-7.4) or M3 medium (15 g L^-1^ soybean oil meal, 15 g L^-1^ glucose, 5 g L^-1^ NaCl, and 1 g L^-1^ CaCO_3_, pH 7.0-7.4) for fermentation (180 rpm) at 30 °C. Production of the target compounds were monitored by UPLC–MS (details in section “UPLC–MS analysis”), and cultures were harvested when production reached a plateau. Culture broths were centrifuged (8000 rpm, 10 min), and supernatants were extracted with XAD-16 resin for 2 h, followed by elution with 50 mL methanol. Eluates were concentrated by rotary evaporation and dissolved in 1 mL of methanol.

### Isotope-labeling precursor feeding experiment

Spores of *S. virginiae* DSM40094*, S. avidinii*-*kasO*p*-*acdA,* and *S. albus* Del14::p15A-*kasO*p\**-nda* for the production of α-methyl-KAPA (**5**), acidomycin (**1**), and ANDA (**6**), respectively, were inoculated into 20 mL of TSB medium and incubated at 30 °C with agitation at 180 rpm for 2-3 days. The resulting seed cultures were then inoculated (1%, v/v) in 10 mL of M6 (20 g L^-1^ D-galactose, 20 g L^-1^ glucose, 2 g L^-1^ (NH_4_)_2_SO_4_, and 2 g L^-1^ CaCO_3_) or M3 media supplemented 1 mM *d*^8^-valine and 1 mM ^13^C_4_^15^N isoleucine to label α-methyl-KAPA, 1 mM ^13^C_5_,^15^N-Cys to label acidomycin, or 0.5 mM ^13^C_5_ glutamic acid to label ANDA. The supernatants were collected on day 1-3 and analyzed by UPLC-MS.

### Metabolite analysis of *E. coli* BW25113

A single colony of *E. coli* BW25113 was inoculated into 1 mL of M9 medium (diluted from 10 × M9 that comprise 60 g L^-1^ Na_2_HPO_4_, 30 g L^-1^ KH_2_PO4, 10 g L^-1^ NH_4_Cl, 5 g L^-1^ NaCl, 2 g L^-1^ glucose, 1 mM MgSO_4_, and 0.1 mM CaCl_2_) containing 1 nM biotin. The overnight culture was centrifuged and the cells were washed with 1 mL of M9 media which did not contain biotin. After the cells were washed twice and resuspended in 1 mL of M9 medium, 1% of the bacterial suspension was transferred to 50 mL of M9 medium. α-Methyl-KAPA at the final concentration of 32 µM was supplemented into the culture after the 4 hours of incubation. After 2 and 16 hours, the metabolites were extracted by adding the same amount of acetone. The extracts were concentrated, diluted with MeOH and applied to UPLC-MS measurement.

### UPLC-MS analysis

The crude extracts were analyzed using UPLC-MS (LC: Ultimate 3000 RS; MS: Bruker amaZon Speed or Bruker maXis4G, Columns: Agilent Zorbax Eclipse Plus C18, 3.5 μm, 4.6 mm × 100 mm; or ACQUITY UPLC BEH C18 Column, 130Å, 1.7 µm, 2.1 mm × 100 mm). Double-distilled water supplemented with 0.1% formic acid and distilled acetonitrile supplemented with 0.1% formic acid were used as eluents. The flow rate of the gradient elution was 0.6 mL min^-1^, and the gradient changed from 5 to 95 % acetonitrile in 18 min and was maintained at 95% acetonitrile for 2 min. The analysis was executed using the Brucker Compass DataAnalysis software (version 5.3).

### Large-scale fermentation and compound isolation

*acidomycin (**1**)*: *S. avidinii-kasO*p*\*-acdA* activation mutant was cultured in 60 300 mL-flasks containing 50 mL of M3 medium. Cultures were incubated at 30 °C, 180 rpm until production reached a plateau and harvested by centrifugation (8000 rpm, 20 min). **1** was separated from the contaminants by liquid-liquid partitioning using with ethyl acetate. The enriched fractions were purified by normal-phase chromatography (Biotage, 200–400 mg crude extract, 25 g silica column; mobile phase A: Hexane, B: ethylacetate, C: MeOH; A/B: 0–100% B, 15 CV; 100% B, 3CV; B/C: 0-100% C, 20CV; 100% C, 5CV). Resulting fractions with targeting compounds were combined and separated using HPLC-MS (LC: Ultimate 3000 RS; MS: Bruker HCT; Column: Kinetex 5 µm Biphenyl 100 Å, LC Column 250 × 4.6 mm). Double-distilled water supplemented with 0.1% formic acid and distilled MeOH supplemented with 0.1% formic acid were used as eluents. The flow rate of the gradient elution was 5 mL min^-1^, and the gradient changed from 5 to 95 % MeOH in 25 min and was maintained at 95% MeOH for 2 min. *α-methyldesthiobiotin (**3**)*: *S. lydicus* wild-type was cultured in five 1 L flasks containing 200 mL of cotton medium (10 g L^-1^ glucose, 10 g L^-1^ dry yeast, 10g L^-1^ cotton meal, and 20g L^-1^ dextrin, dissolved in tap water) and 10 days later the supernatant was collected by centrifugation of the culture broth. The supernatant was applied to the column packed with 100 mL of the activated carbon with 20-60 mesh and **3** was eluted with 0-10% of acetone stepwise. The fractions containing **3** was collected and further purified by liquid–liquid partitioning with *n*-butanol. The enriched fractions were purified by reversed-phase chromatography (Biotage, 25 g C18 column; mobile phase A: H₂O, B: ACN) was repeated with the LC conditions of gradient mode with 5-95% of B and isocratic mode with 10% of B. Resulting fractions were combined and compounds were separated using HPLC-MS (LC: Ultimate 3000 RS; MS: Bruker HCT; Column: Agilent Zorbax Eclipse Plus C18). 84% of double-distilled water supplemented with 0.1% formic acid and 16% of distilled acetonitrile supplemented with 0.1% formic acid were used as eluents in isocratic mode with the flow rate of 5 mL min^-1^. In addition, the semi-preparative purification with the Phenyl column (XBridge® BEH Phenyl OBD^TM^ Prep Column, 130 Å, 5 µm, 10 mm × 250 mm) was carried out under the LC condition of 5-95% of B in 20 min. The obtained **3** was dried with an evaporator and with N_2_ gas, which was subsequently applied to NMR measurement.

*α-methyl-KAPA (**5**)*: *S. virginiae* wild-type was cultured in five 1L-flasks containing 200 mL of M6 medium and five days later, the supernatant was obtained by centrifugation of the culture broth. The supernatant was applied to the column packed with 100 mL of the activated carbon with 20-60 mesh and **5** was eluted with 0-30% of acetone stepwise. The fractions containing **5** were collected and the solvent was removed by an evaporator. The residue was dissolved in water and applied to the column packed with 100 mL of weak acid cation exchange (MAC3H) and the compound was eluted with 0-0.2 N formic acid stepwise. The enriched fractions were purified by reversed-phase chromatography (Biotage, 25 g C18 column; mobile phase A: H₂O, B: ACN) was repeated with the LC conditions of 2% of B isoclatic mode and 5% of B isoclatic mode. Resulting fractions were combined and compounds were separated using HPLC-MS (LC: Ultimate 3000 RS; MS: Bruker HCT; Column: Agilent Zorbax Eclipse Plus C18). 98% of double-distilled water supplemented with 0.1% formic acid and 2% of distilled acetonitrile supplemented with 0.1% formic acid were used as eluents in isocratic mode with the flow rate of 5 mL min^-1^. The solvent of the fractions containing **5** was removed by an evaporator and the compound was immediately dissolved in D_2_O to apply to NMR analysis to avoid the spontaneous dimerization.

*ANDAs (**6,*** ***6*, 10****), stravidin* ***8****, and compound* ***9***: *S. albus* Del14::p15A-*kasO*p*-*nda* (ANDAs), *S. avidinii* DSM40526 (stravidin **8**), and *S. albus* Del14::p15A-*kasO*p*-*mkp* (compound **9**) was inoculated (10%, v/v) with 3-day seed cultures into 5 L flask containing M3 media. Cultures were incubated at 30 °C, 180 rpm until production reached a plateau and harvested by centrifugation (8000 rpm, 20 min). Supernatants containing **8** and **9** were extracted with 2% (v/v) XAD-16N resin, whereas those containing **6**, **6\*** and **10** were extracted with 3% (w/v) activated charcoal, both stirred overnight. The resulting adsorbents were collected, lyophilized, and sequentially extracted (each 2 L, containing 0.1% formic acid) with hexane, ethyl acetate, acetone, and methanol using a peristaltic pump. Cell pellets were extracted in 1 L of methanol overnight. Organic extracts containing the targeted compounds were concentrated under reduced pressure and redissolved in methanol (**6**, **6\*** and **10** in H₂O), followed by liquid–liquid partitioning with *n*-butanol. The enriched fractions were purified by reversed-phase chromatography (Biotage, 200–400 mg crude extract, 25 g C18 column; mobile phase A: H₂O + 10 mM ammonium formate, B: MeOH + 10 mM ammonium formate; 0–40% B, 30 CV; 40–100% B, 10 CV). Resulting fractions with targeting compounds were combined and separated using HPLC-MS (LC: Ultimate 3000 RS; MS: Bruker HCT; Column: Agilent Zorbax Eclipse Plus C18). Double-distilled water supplemented with 0.1% formic acid and distilled MeOH supplemented with 0.1% formic acid were used as eluents. The flow rate of the gradient elution was 5 mL min^-1^, and the gradient changed from 5 to 95 % MeOH in 25 min and was maintained at 95% MeOH for 2 min.

### Chemical modification of Ala-ANDA (10)

To chemically modify Ala-ANDA (**10**), the compound was dissolved in methanol and subjected to ozonolysis. The resulting aldehyde intermediate was subsequently oxidized with 30% aqueous hydrogen peroxide to afford an Ala–Glu dipeptide for subsequent Marfey’s method.

### Advanced Marfey’s method

The compound was hydrolyzed in 100 μL of 6N HCl at 110 °C for 1 hour, followed by drying under nitrogen stream. The resulting hydrolysate was dissolved in 110 μL of H_2_O and divided into two equal portions (50 μL each). To each aliquot, 20 μL of 1N NaHCO_3_ and 20 μL of 1% Marfey’s reagent (either L-FDLA or D-FDLA) were added sequentially. The reaction mixtures were incubated at 45 °C with shaking at 700 rpm for 2 hours. Each reaction was then neutralized with 10 μL of 2N HCl and diluted with 300 μL of acetonitrile. Standard amino acids were derivatized under the identical conditions. The derivatized samples were centrifuged and analyzed by UHPLC-MS using an ACQUITY BEH column (100 × 2.1 mm 1.7 μm 130 Å) at a flow rate of 0.6 mL min^-1^ and a column temperature of 45 °C. Eluent A consists of H_2_O with 0.1% formic acid, and eluent B of acetonitrile with 0.1% formic acid. The gradient was as follows: 0-1 min, 5% to 10% eluent B; 1-15 min, 10% to 35% eluent B; 15-22 min, 35% to 55% eluent B; 22-25 min, 55% to 95% eluent B. Mass spectrometric detection was performed in positive ion mode.

### Structure elucidation

*acidomycin (**1**)*: Compound **1** was assigned the molecular formula by HRESIMS data (*m/z* 218.0844 [M+H]^+^). 1D and HSQC NMR spectra (Supplementary Figs. 67-71, Supplementary Table 14) indicated five methylenes, one methines, and two quaternary carbons. Detailed analysis of COSY and HMBC spectra revealed **1** is the known antibiotic acidomycin^38^. The optical rotation value of **1** in MeOH was [α]_D_^20^ −44°, which is consistent with the reported value of [α]_D_^23^ −46.0° (c 1.00, MeOH) for (*S*)-(−)-acidomycin^38^.

*α-methyldesthiobiotin (**3**)*: Compound **3** was assigned the molecular formula C_11_H_20_N_2_O_3_ by HRESIMS data (*m/z* 229.1550 [M+H]^+^). 1D and HSQC NMR spectra (Supplementary Figs. 43-47, Supplementary Table 9) indicated two methyl, four methylenes, three methines, and two quaternary carbons. Further analysis of the COSY and HMBC spectra revealed **3** is the known compound α-methyldesthiobiotin^17,18^ *α-methyl-KAPA (**5**)*: Compound **5** was assigned the molecular formula C_10_H_19_NO_3_ by HRESIMS data (*m/z* 202.1438 [M+H]^+^). 1D and HSQC NMR spectra (Supplementary Figs. 33-37, Supplementary Table 7) indicated two methyl, four methylenes, two methines, and two quaternary carbons. Detailed analysis of COSY and HMBC spectra revealed that **5** is the known compound 8-amino-2-methyl-7-oxononanoic (α-methyl-KAPA).^20^

*ANDA (**6**)*: Compound **6** was assigned the molecular formula C_9_H_13_NO_4_ by HRESIMS data (*m/z* 200.0920 [M+H]^+^). 1D and HSQC NMR spectra (Supplementary Figs. 48-51, Supplementary Table 10) indicated two methylenes and five methines. Analysis of the 2D NMR data revealed **6** as 2-aminonona-5,7-dienedioic acid, a subunit present in compound **8**. The configurations of two double bonds were determined to be 5*E*,7*E* based on their large coupling constants.

*(E)-**6\****: Compound (*E*)-**6\*** was assigned the molecular formula C_9_H_13_NO_4_ by HRESIMS data (*m/z* 200.0917 [M+H]^+^). The NMR spectra of (*E*)-**6\*** were similar to that of **6** (Supplementary Figs. 57-61, Supplementary Table 12). Careful analysis indicated (*E*)-**6\*** is a cyclized derivative of **6**, evidenced by the disappearance of one double bond, downfield shift of C-5 (61.7 ppm), and the HMBC correlation from H-2 (4.15 ppm) to C-5 (61.7 ppm). The configuration of the double bond was determined to be 6*E* based on the large coupling constant. Moreover, (*E*)-**6\*** is speculated to be a mixture of stereoisomers indicated by the two sets of NMR signals observed, with the difference at the absolute configuration of C-5.

*(Z)-**6\****: Compound (*Z*)-**6\*** was assigned the molecular formula C_9_H_13_NO_4_ by HRESIMS data (*m/z* 200.0923 [M+H]^+^). The NMR spectra of (*Z*)-**6\*** were almost identical to those of (*E*)-**6\*** (Supplementary Figs. 62-66, Supplementary Table 13). Careful analysis of the NMR data showed the only difference as the configuration of the double bond, 6*Z* in (*Z*)-**6\*** while 6*E* in (*E*)-**6\***, respectively.

*Ala-ANDA (**10**)*: Compound **10** was assigned the molecular formula C_12_H_18_N_2_O_5_ by HRESIMS data (*m/z* 271.1288 [M+H]^+^). Analysis of the 1D and HSQC NMR spectra (Supplementary Figs. 52-56, Supplementary Table 11) revealed one methyl, two methylenes, six methines (four olefinic), and three quaternary carbons. Detailed interpretation of the 1D and 2D NMR data indicated the presence of one alanine residue and ANDA residue. The connectivity between these two units was established based on the HMBC correlation from H-2 (4.29 ppm) to C-1’ (170.4). Finally, the absolute configurations at C-2 and C-2’ were determined as 2*S*,2’*S* through chemical modifications followed by Marfey’s method as mentioned above.

*stravidin* ***8***: Compound **8** was assigned the molecular formula C_19_H_31_N_3_O_4_ by HRESIMS data (*m*/*z* 366.2384 [M+H]^+^). Analysis of the 1D and HSQC NMR spectra (Supplementary Figs. 23-32, Supplementary Table 6) revealed four methyls, three methylenes and nine methines. Analysis of 2D NMR data revealed one *N*-methylleucine residue and one *N*-acetyl-amiclenomycin (AcAcm) residue. Although no direct HMBC correlations were observed, the connectivity between these two units was inferred based on the chemical shifts of the α-methine protons of Leu (3.51 ppm) and AcAcm (4.30 ppm).

*Compound* ***9***: Compound **9** was assigned the molecular formula C_20_H_32_N_2_O_4_ by HRESIMS data (*m/z* 365.2444 [M+H]^+^). Examination of the 1D and HSQC NMR spectra (Supplementary Figs. 38-42, Supplementary Table 8) revealed two methyl, four methylenes, one methines, and three quaternary carbons. Further analysis of the COSY and HMBC correlations established the connectivity of all carbon atoms. In combination with the molecular formula, these data suggested that **9** possesses a symmetric planar structure. Accordingly, the planar structure of **9** was determined to be a tetrasubstituted pyrazine. The absolute configurations at C-2 and C-2’ were proposed to be 2*R*,2’*R* by comparison of the optical rotation of **9**, [α]_D_^22^ −14.6° (c 0.3, CHCl_3_), with those of (*S*)-2-methylhexanoic acid, [α]_D_^20^ +20.6° (c 0.5, CHCl_3_), and (*R*)-2-methylhexanoic acid, [α]_D_^22^ −18° (c 1.1, CHCl_3_), respectively.

### Protein overexpression and purification

pHisTev-*sav1*, pHisTev-*sav2*, pET28-*bioA*, pET28-*bioB*, pColdI-*mdbA*, and pColdI-*mdbB* were constructed and electroporated into *E. coli* BL21(DE3) for protein expression.

The *E. coli* BL21(DE3) carrying the corresponding plasmid was cultured in LB or 2 × YT medium supplemented with respective antibiotics at 37°C overnight. 1% of an overnight culture was inoculated into large-scale LB or 2 × YT medium containing respective antibiotics and incubated at 37°C until OD_600_ reached 0.4-0.6. Protein expression was induced by adding isopropyl β-D-1-thiogalactopyranoside (IPTG) to a final concentration of 0.1 mM after cooling into 16°C and the culture was further incubated at 16°C overnight. The cells were harvested (8000 rpm, 4°C, 15 min) and resuspended in lysis buffer (20 mM Tris pH8.0, 200 mM NaCl, 10%(v/v) glycerol). After the cell disruption using with a sonicater or French press, the cell lysis was centrifuged at 40,000 × *g* for 30 min at 4 °C and applied to Ni-NTA agarose column connected to Äkta Go purifier (Cytiva). After washing the column with washing buffer (lysis buffer + 20 mM imidazole pH8.0), protein was eluted using elution buffer (lysis buffer + 250 mM imidazole pH 8.0). For the purification of the recombinant Sav1, PBS buffer pH 8.0 was used instead of Tris. Protein was desalted using PD-10 Desalting Column (GE) and eluted with lysis buffer. Protein purity was confirmed by SDS-PAGE or Maxis 4G. Protein was concentrated by Amicon® Ultra Centrifugal Filters (Millipore) and stored at −80°C after the fast-frozen by liquid N_2_. Streptavidins were further desalted by Amicon® Ultra Centrifugal Filter 0.5 mL (Millipore) with PBS buffer, pH 7.0, directly before the use for the bioassay. Protein concentration was determined spectrophotometrically using calculated extinction coefficient from the amino acid sequence using the PROTPARAM webserver (http://web.expasy.org/protparam/)^45^ or by the Bradford method (Bio-Rad Protein Assay Dye Reagent Concentrate).

### *In vitro* BioAD coupling assay with α-methyl-KAPA (5)

*E. coli* derived BioA and BioB were employed for the assays. Reactions (50 µL) contained 100 mM HEPES (pH 8.5), 5 mM SAM, 3.125 µM 7-keto-8-aminopelargonic acid (KAPA), 50 mM NaHCO₃, 5 mM ATP, 1 mM MgCl₂, 1 mM pyridoxal 5’-phosphate (PLP), 1 µM BioA, 2 µM BioD, and 32 µM testing compounds such as α-methyl-KAPA (**5**). Control reactions omitted individual components as indicated. After incubation for 16 h at 30 °C, reactions were quenched with 50 µL methanol and centrifuged (21,500 × *g*, 10 min). Supernatants were analyzed as described above, and mass spectra were acquired on an amaZon Speed instrument.

### *In vitro* conversion of α-methyldesthiobiotin (3) using MdbAB from α-methyl-KAPA (5)

For the assay of MdbAB (the second copy of BioAD in *S. lydicus*), 100 µL reaction mixture was prepared in 1.5 mL Eppendorf tubes, containing 14 μM of MdbA, 25 μM of MdbB, 0.5 mM SAM, 20 µM PLP, 1 mM ATP, 1 mM MgCl_2_, 2 mM NaHCO_3_ and partially purified **5** incubated at 30 °C. After the overnight reaction, the mixture was measured by UPLC-MS system.

### Bioactivity assay using *E. coli* BW25113

A single colony of *E. coli* BW25113 was inoculated into 1 mL of M9 medium containing 1 nM biotin. The overnight culture was centrifuged and the cells were washed with 1 mL of M9 medium which do not contain biotin. After the cells were washed twice and resuspended in 1 mL of M9 medium, 50 µL of the bacterial suspension was transferred to 5 mL of M9 medium. The OD_600_ was measured four hours later and bacterial suspension in M9 medium was prepared so that the cell concentration was around 1.0 × 10^6^ CFU mL^-1^. 50 μL of the bacterial suspension per well was dispensed into round-bottom clear sterile polypropylene plates (Costar) containing the testing compounds or spectinomycin as a positive control^46^ serially diluted between 256 to 0.13 μM at the final concentration after mixing with the bacterial suspension, depending on the compound in the same medium at 50 μL per well. Plates were shaken at 750 rpm for 19 hours at 37 °C followed by the addition of 20 μL of 0.02 % resazurine solution and the incubation for 30 min under the same condition. Cell viability was calculated based on the absorbance at 570 nm, measured with a SpectraMax T5 plate reader (Molecular Devices). KAPA or DAPA was added at the final concentration of 0.1 μM or 1 μM, respectively, into the well containing *E. coli* Keio mutant Δ*bioF* and 2.1 μM of α-methyl-KAPA. 131 µM streptavidin 1 in PBS, pH 7.0, was treated in a well containing 64 µM acidomycin at RT before the bacterial suspension was dispensed. All assays were performed in biological replicates, and error bars represent the SEM. All statistical analyses were conducted using one-way ANOVA with multiple comparisons. Bar chart in Fig. 5d was generated using GraphPad Prism 10.5.

### Protein expression and purification of Sav2 for crystallization

The DNA fragment encoding Sav2 (WP_189973542.1, residues 45–169) was amplified by PCR and cloned into the pET28a (+) vector using TEDA assembly^47^. The protein expression plasmid was transformed into *E. coli* BL21 (DE3). A single colony was inoculated into a culture medium and grown overnight. Subsequently, 10 mL of this overnight culture was used to inoculate 2 L of LB medium and grown at 37 °C with shaking at 180 rpm for 4 hours. After pre-cooling at 16 °C for 2 hours, protein expression was induced by adding 0.1 mM IPTG and continued overnight under the same conditions. Cells expressing Sav2 were harvested and resuspended in 70 ml of binding buffer (20 mM Tris-HCl, 500 mM NaCl, 20 mM imidazole, pH 8.0), followed by lysis via sonication. The lysate was centrifuged at 15,000 × *g* for 1 hour to remove debris, and the supernatant was loaded onto a Ni-NTA agarose column (Bestchrom). The N-terminal hexa-histidine-tagged protein was cleaved with 100 U of thrombin (Solarbio) for 12 hours at 4 °C. Finally, the tag-free protein was further purified using a Superdex 200 pg gel-filtration column (Cytiva) equilibrated with a buffer containing 20 mM Tris and 400 mM NaCl, pH 8.0.

### Crystallization of Sav2-acidomycin (1) and X-ray data collection

Sav2 (20 mg mL^-1^) was incubated with **1** (6 mM) on ice for 1.5 hours. Crystallization of the Sav2-**1** complex was performed at 16 °C using the hanging-drop vapor diffusion method by mixing equal volumes of the protein complex and reservoir solution. The reservoir solution contained 15% (v/v) 2-propanol, 0.1 M MES monohydrate pH 6.0, and 18% (w/v) polyethylene glycol monomethyl ether 2,000. 25% glycol was added into the reservoir solution to cryoprotect the crystals before flash frozen. Diffraction data were collected at 100 K on beamlines BL18U1 at the Shanghai Synchrotron Radiation Facility (SSRF)^48^, and processed using the HKL3000 program^49^.

### Structure determination and refinement

The structure of Sav2-**1** was solved by molecular replacement with the program Phaser^50^, using the sreptavidin structure (PDB code: 1STP^22^) as the search model. Further manual model building was facilitated by using Coot^51^, combined with the structure refinement using Phenix^52^. Data collection and structure refinement statistics are summarized in Supplementary Table 5. The Ramachandran statistics, as calculated by Molprobity^53^, are 97.5%/0 (favored/outliers). All the structural figures were prepared using PyMol (Schrödinger, LLC).

## Data availability

The protein crystal structure data generated in this study have been deposited in the Protein Data Bank under PDB ID 9XIK. The biosynthetic gene cluster sequences in this study are provided as supplementary data in the supplementary information. All data that support the findings of this study are available in the main text, supplementary information and from corresponding author(s) upon request. Source data are provided with this paper.

## Acknowledgements

This work was supported by the Helmholtz International Lab (InterLabs-0007). Research in the laboratories of Chengzhang Fu and Rolf Müller was funded by the Bundesministerium für Bildung und Forschung (BMBF). Sumire Kurosawa was partially financed by the Naito Foundation and the Uehara Memorial Foundation. The authors thank Alexander Popoff for assistance with structure elucidation and Vidhisha Sonawane for help with bioassays. The authors also thank Xueyun Geng from Core Facilities for Life and Environmental Sciences of Shandong University for her assistance during X-ray diffraction data collection, and the staff members of BL18U1 (https://cstr.cn/31129.02.NFPS.BL18U1) at the National Facility for Protein Science in Shanghai for providing technical support and assistance in data collection and analysis.

## Contributions

C.F. conceived and supervised the study. C.F. and S.K. analyzed the gene clusters. C.F., S.K., J.M., F.X., and T.W. performed ACTIMOT activation, heterologous expression of BGCs and *in vitro* biochemical studies. S.K. performed the bioactivity assays. X.D., J.M., and D.W. performed the protein crystallographic studies. S.K., J.M., and T.W. isolated the compounds, S.K. and H.Z. determined the chemical structures of compounds. C.F., S.K., J.M., F.X., X.D., D.W., H.Z. and R.M. analyzed the data. C.F. and R.M. raised the funding for this study. C.F., R.M., S.K., J.M., and F.X. wrote the paper with input from all authors. S.K., J.M., X.D., and F.X. contributed equally to the manuscript. All authors read and approved the final manuscript.

